# Insights into the role of root exudates in bacteriophage infection dynamics

**DOI:** 10.1101/2024.07.15.603626

**Authors:** Vlastimil Novak, Michelle C. M. van Winden, Thomas V. Harwood, Rachel Neurath, Suzanne M. Kosina, Katherine B. Louie, Matthew B. Sullivan, Simon Roux, Karsten Zengler, Vivek K. Mutalik, Trent R. Northen

## Abstract

Bacteriophages impact soil bacteria through lysis, altering the availability of organic carbon and plant nutrients. However, the magnitude of nutrient uptake by plants from lysed bacteria remains unknown, partly because this process is challenging to investigate in the field. In this study, we extend ecosystem fabrication (EcoFAB 2.0) approaches to study plant-bacteria-phage interactions by comparing the impact of phage-lysed and uninfected ^15^N-labeled bacterial necromass on plant nitrogen acquisition and rhizosphere exometabolites composition. We show that grass *Brachypodium distachyon* derives some nitrogen from amino acids in uninfected *Pseudomonas putida* necromass but not from virocell necromass. Additionally, the bacterial necromass elicits the formation of rhizosphere exometabolites, some of which (guanosine), alongside tested aromatic acids (*p*-coumaric and benzoic acid), show distinct effects on bacteriophage-induced lysis when tested *in vitro*. The study highlights the dynamic feedback between bacterial necromass and plants and suggests that root exudate metabolites can impact bacteriophage infection dynamics.

## 1. Introduction

It is well established in marine environments that predation by bacteriophage (phage), viruses that infect and replicate within bacterial cells, can have ecologically important impacts on the infected host ‘virocell’ metabolism (Clokie, Millard, Letarov & Heaphy 2011; Howard-Varona *et al*. 2020). Furthermore, the ‘viral shunt and shuttle’ of released nutrients from the phage-lysed bacterial cells can significantly contribute to marine nitrogen (N) and carbon (C) cycling (Gobler, Hutchins, Fisher, Cosper & Saňudo-Wilhelmy 1997; Breitbart, Bonnain, Malki & Sawaya 2018). In contrast, soil ecosystems, known to contain abundant and diverse phages, lack such functional characterization (Emerson 2019; Bi *et al*. 2021; Roux & Emerson 2022). This is primarily due to the soil’s complexity and the heterogeneity of ionic strength necessary for phage infectivity (Kuzyakov & Razavi 2019; Carlson *et al*. 2023).

Recent evidence highlighted the re-utilization of viral lysate-derived nutrients by surviving microbes and the release of recalcitrant C as evidence for a viral shunt and shuttle in soil, respectively (Tong *et al*. 2023). Phages were also recently used to manipulate soil bacterial communities, affecting N availability (Braga *et al*. 2020). The phage lysis of rhizosphere bacteria was shown to redistribute plant-derived C into the environment (Starr *et al*. 2021). Nevertheless, the impact of phages on rhizosphere N cycling, and subsequently on plant growth, remains poorly understood despite the potential of phages to increase N availability in forms readily absorbed by roots, including ammonium (NH_4_^+^), nitrate (NO_3_^−^) and organic molecules like amino acids (Näsholm, Kielland & Ganeteg 2009; Muratore, Espen & Prinsi 2021).

Ecosystem fabrication (EcoFAB) is a laboratory-based approach to studying host and microbial interactions under highly controlled conditions (Zengler *et al*. 2019). EcoFAB has been used to study root exudation dynamics, synthetic community assembly processes, non-invasive plant phenotyping, and high-resolution root imaging (Sasse *et al*. 2019; Jabusch *et al*. 2021; Kuang *et al*. 2022; Acharya *et al*. 2023; Lin *et al*. 2023). Recently, we used improved EcoFAB 2.0 devices to demonstrate the impact of inorganic N sources on the abundance of N-containing compounds in root exudates (Novak *et al*. 2024).

Our objective was to use EcoFAB 2.0 to study the effects of virocell necromass on plant N acquisition. We hypothesized that phage-lysed bacteria can provide nitrogen for plant uptake. To test this hypothesis, we supplied ^15^N isotopically labeled viral lysates (using lytic gh-1 phage) and sonicated lysates of the model rhizosphere bacterium *Pseudomonas putida* to the model grass *Brachypodium distachyon*, grown hydroponically in EcoFABs 2.0 (Lee & Boezi 1966; Liffourrena & Lucchesi 2018). After one week of incubation with plants, we analyzed the medium for changes in metabolite composition by LC-MS/MS. We also traced the uptake of bacterial N into the plant through stable isotope labeling. Follow-up bioassays evaluated the impact of specific root exudate compounds, elicited by necromass, on the growth of two rhizosphere-dwelling bacteria infected with lytic phages, *P. putida* with phage gh-1 and *B. subtilis* with phage SPO1 (Okubo, Strauss & Stodolsky 1964; Gallegos-Monterrosa, Mhatre & Kovács 2016), revealing distinct effects of plant-derived metabolites on phage lysis *in vitro*. This study suggests that plant exudates can impact bacteriophage predation in the rhizosphere.

## 2. Results

### 2.1. The effect of lysed bacteria on plant biomass

First, we tested the effect of *P. putida* A.3.12 on the growth of *B. distachyon* by measuring root and shoot biomass, indicating no inhibition of inoculated plants compared to the axenic control (**Fig. S1**). Consequently, we proceeded with all subsequent experiments using *P. putida* cultures grown on unlabeled or ^15^N-labeled substrates to create necromass media for plants. The media were prepared by sterile-filtering (pore size 0.2 µm) of virocells (i.e., phage-lysed), sonicated cells, and unlysed cells diluted to 0.1x concentration in an N-free medium, subsequently supplied to plants for one week (**Fig. 1a**). Our controls were axenic plants grown on an N-free medium and a technical control without plants. To estimate lysis, we enumerated cells by microscopy in undiluted unlabeled cultures (**Fig. S2a)**. Compared to unlysed culture with 3.8e9 cells/ml, phage exposure decreased cell density by 61%, while sonication resulted in 88% lysis. For conversions between microscopy enumeration and optical densities of *P. putida*, we established a calibration curve: cells/ml=2.281e9*OD (**Fig. S2b**).

**Fig. 1:**
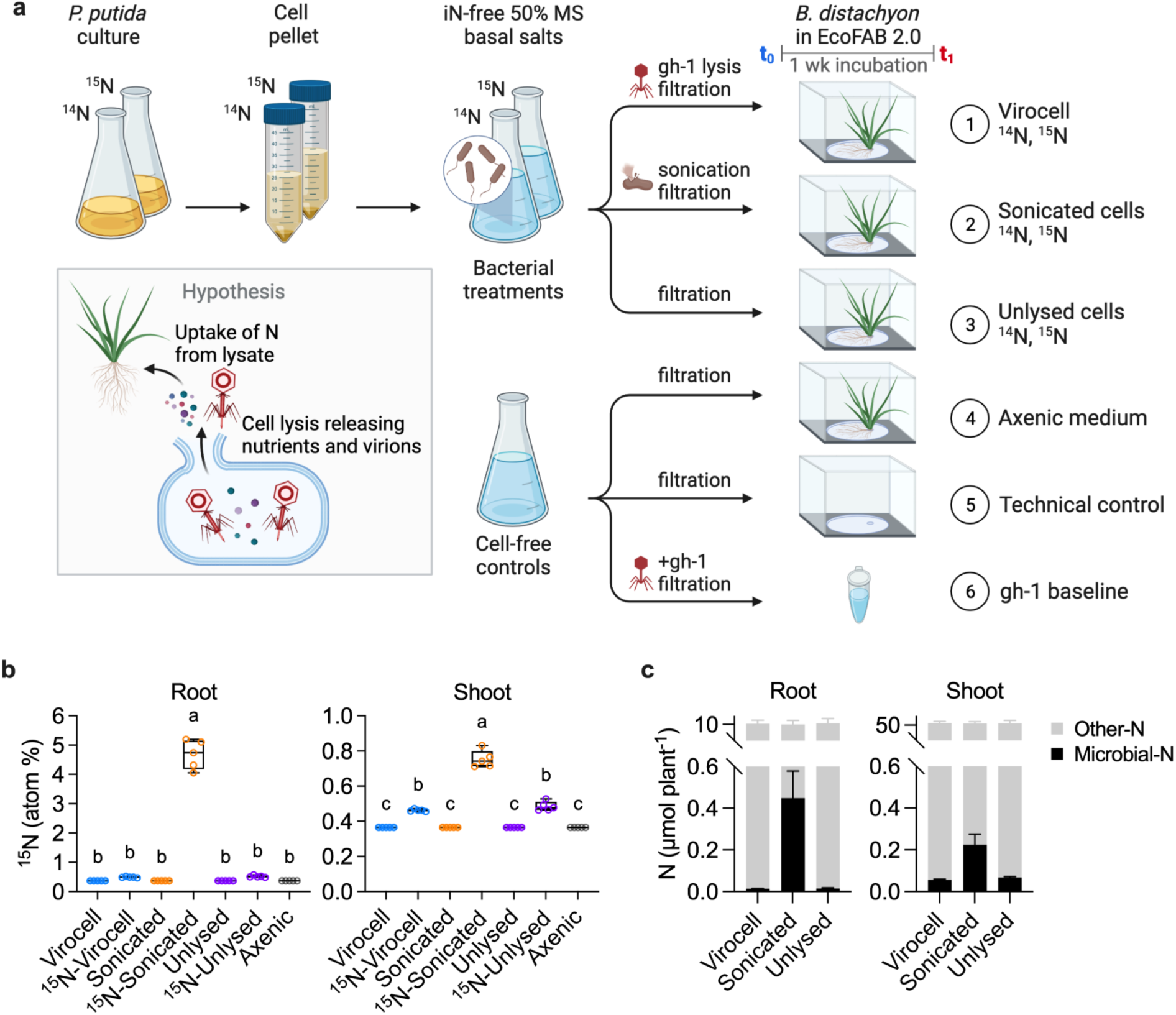
Impact of bacterial necromass on plant nitrogen. (**a**) Experimental design: ^15^N-labeled and unlabeled bacterial lysates were supplied to plants for one week to test the hypothesis that bacterial necromass provides N for plant uptake. (**b**) Stable isotope analysis revealed ^15^N-enrichments in labeled treatments. Box plots display all data points, with hinges spanning the 25th to 75th percentiles, a central line denoting the median, and whiskers reaching the minimum and maximum values. Different letters indicate significant differences at *p*<0.05, One-way ANOVA with Tukey’s test, *n*=5. (**c**) Microbial-N contribution to total plant N in roots and shoots was the biggest from the sonicated cells. Values are mean±SD.

To study the effects of bacterial necromass on the plant, we used the ^15^N-labeled and unlabeled necromass solutions to incubate 3-week-old *Brachypodium* plants in EcoFAB 2.0 for 1 week (**Fig. S3a**), followed by shoot and root biomass analysis. Necromass, generated by phage lysis or sonication, caused no significant changes in plant fresh or dry weights (**Fig. S3b**) or the total plant N content (**Fig. S3c**). However, stable isotope analysis showed significant ^15^N-enrichment in plants supplied with labeled necromass treatments (**Fig. 1b**). Plant shoots contained 5 times more total N than roots, with small (0.1–4.4%) microbial-N contribution, which was the highest in roots supplied with sonicated but not with the viral necromass (**Fig. 1c**).

To evaluate if plants depleted N and C from the necromass medium, we measured total nitrogen (TN) and dissolved organic carbon (DOC) concentrations, which showed a general decrease to the medium baseline after incubation with the plants (**Fig. S3d**). While *P. putida* (alongside *P. simiae*) cells utilize NH_4_^+^, their lysis generally releases negligible amounts of inorganic N relative to protein (**Fig. S4**). Based on this, the N requirements of 4-week-old *Brachypodium* could be theoretically satisfied by >3.3e10 lysed *Pseudomonas* cells (**Table S2**), which is 10 times more than the lysed cell quantity supplied in sonicated treatments.

We investigated the effects of prolonged necromass exposure on plant growth. *B. distachyon* grew for 3 weeks with two dilutions of sonicated bacterial necromass (0.1x or 0.2x lysate in N-free medium). The increasing lysate concentration did not affect shoots or root biomass relative to control plants without necromass (**Fig. S5a**). Notably, a precipitate formed in the rhizosphere of plants supplied with 0.2x lysate (**Fig. S5b**).

### 2.2. Microbial necromass modulates metabolite composition in the rhizosphere

Since virocells are known to have altered metabolism relative to uninfected cells and plants can take up and produce metabolites, we used LC-MS/MS analysis (**Table S1**) for a metabolomics comparison of necromass media before (t_0_) and after (t_1_) a one-week incubation with plants. The GNPS was then used for feature annotation against spectral libraries, resulting in 2059 features (417 annotated) that were used for untargeted metabolomics (**Table S3**). We detected the highest number of unique features in the virocell at t_0_ (**Fig. 2a**) and many features unique to sonicated cells that were absent in the virocells. The virocell exometabolome at t0 results from a 1-hour incubation with phage. Comparing the virocell medium to the gh-1 phage inoculum solution, we observed differences in features abundance (**Fig. S6a**), molecular mass (**Fig. S6b**), and a decrease of amino acids/peptides in the virocell medium (**Fig. S6c**).

**Fig. 2:**
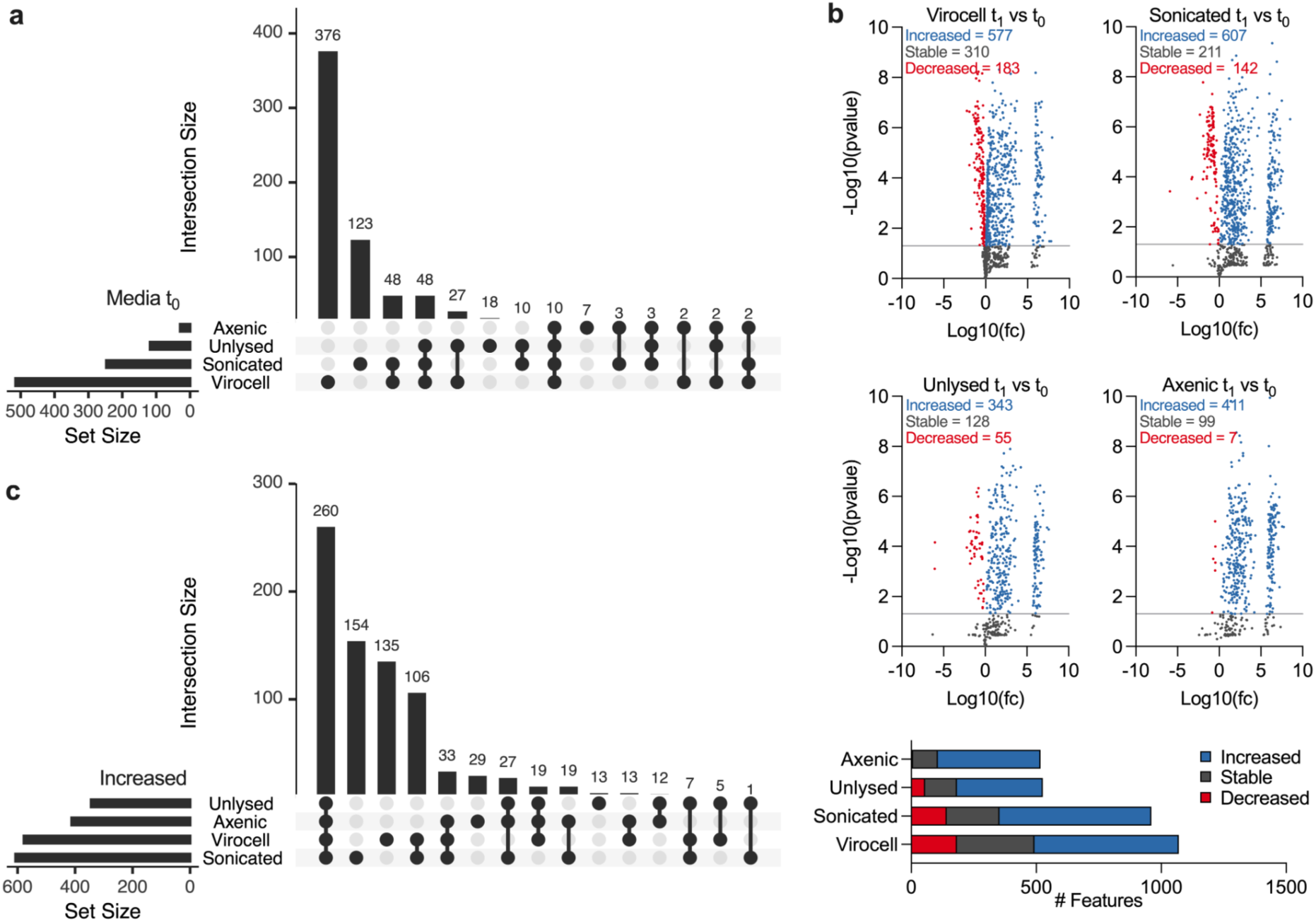
Untargeted metabolomics for bacterial necromass transformations in the rhizosphere of *Brachypodium distachyon*. (**a**) The UpSet plot shows shared features with max peak height >1e6 across necromass media at t_0_. (**b**) Volcano plot for peak heights of metabolomic features showing fold-change (fc) and *p*-value between t_1_ and t_0_. Points above the gray line are statistically significant (*p*<0.05, t-test, *n*=5), indicating stable (gray), up-regulated (blue), and down-regulated (red) features during plant incubation. We included only features with at least one peak height >1e6 across all samples. (**c**) Intersecting Up-regulated features.

We compared metabolite profiles between time points (t_0_ vs t_1_) to assess metabolites that decreased (turnover via plant uptake or degradation) or increased (chemical transformations or root exudation) during incubation with plants (**Fig. 2b**). Larger numbers of features significantly decreased in intensity for viral and sonicated necromass (183 and 142 features, respectively), compared to features in plant incubations with unlysed cells (55) or axenic controls (7). In all treatments, there was a consistently large number of features (343-607) where the intensity increased in the rhizosphere after one week, potentially as a result of root exudation (**Fig. 2b**); the majority of these features (260) were shared between virocell, soniced, axenic, and unlysed treatments (**Fig. 2c**). Unexpectedly, viral or sonicated necromass caused an increase of several unique features (**Fig. 2c**).

To trace metabolite origin, we analyzed the metabolomics data from labeled treatments to determine the relative unlabeled (M0) and ^15^N-labeled (M1+n) intensities of each metabolite (**Fig. 3**). Generally, higher metabolite ^15^N-labeling was present in virocell vs. sonicated treatments. Several amino acids (glycine, tryptophan, glutamic acid, allothreonine, threonine, alanine, serine, and valine) from sonicated necromass decreased in M1+n intensity, indicating the plant took up this microbial-derived N. Notably, *B. distachyon* produced nucleoside intensity (e.g., cytidine, guanosine) with high levels of ^15^N-labeling, indicating their formation from microbial N.

**Fig. 3:**
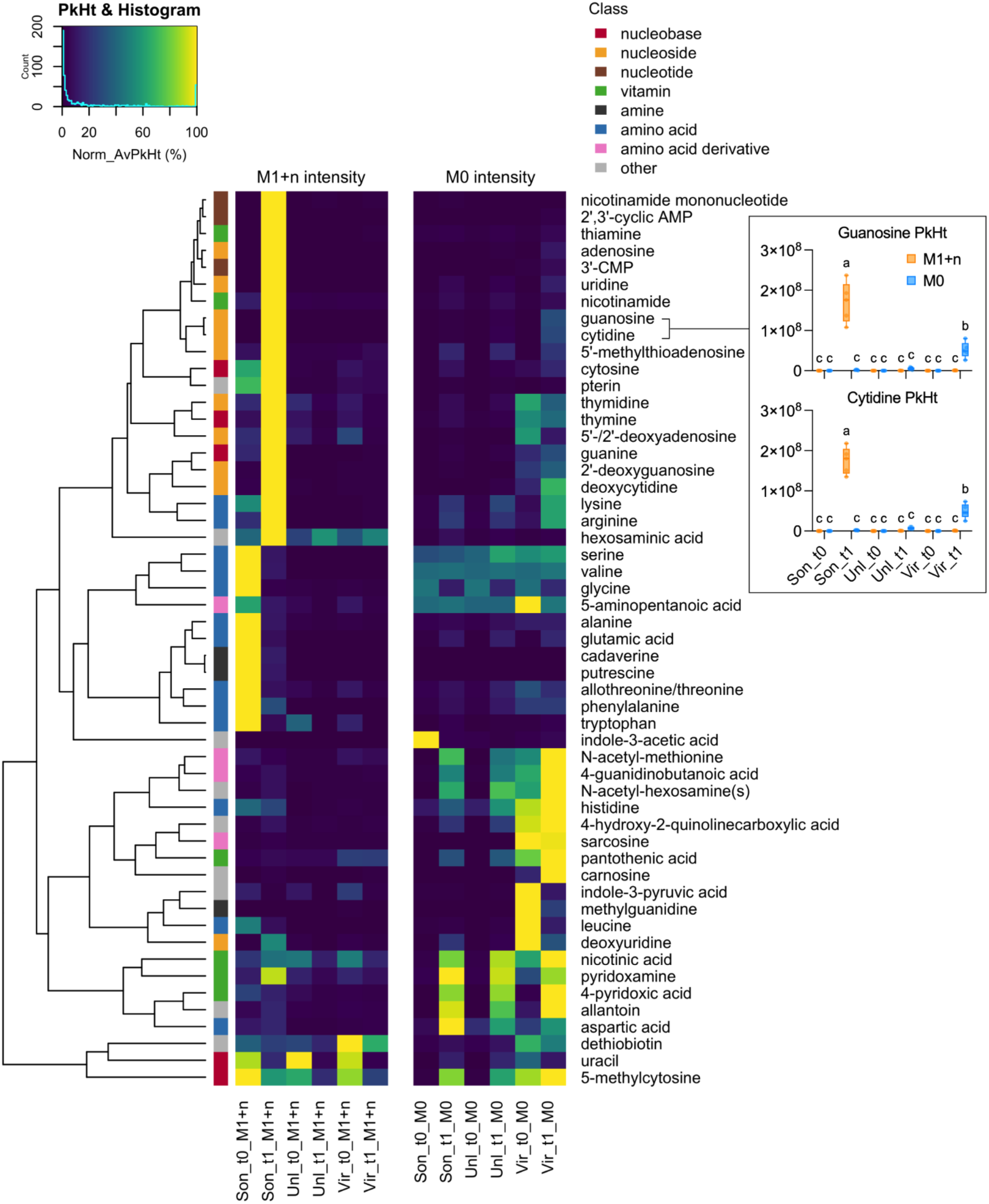
Stable isotope tracing reveals metabolite origin during the transformation of ^15^N-necromass in the rhizosphere. The heatmap shows row normalized labeled (M1+n) and unlabeled (M0) isotopologue peak height for N-containing metabolites in ^15^N-labelled treatments during one week of incubation (t0 vs t1) of bacterial (virocell, sonicated, or unlysed) necromass in *B. distachyon* rhizosphere. The t_0_ time point represents the necromass solution before addition to the plant. Box plots show isotopologue intensities for guanosine and cytidine, displaying all data points, 25th to 75th percentiles, median, and min–max values. Different letters indicate significant differences at *p*<0.05, Two-way ANOVA with Tukey’s test, *n*=5.

Next, we performed a follow-up necromass turnover experiment to determine if plants mediate nucleoside formation (**Fig. S7**). Our experiment demonstrated that while some amino acids result from necromass turnover without plants, *B. distachyon* facilitates the production of most N compounds, including the nucleosides cytidine and guanosine.

### 2.3. Exometabolites modulate phage lysis dynamics

The observation that virocell necromass elicited the production of specific molecules in the rhizosphere suggested that these metabolites may have some phage-related activities (**Fig. 3**). To examine this, we tested the effect of nine diverse root exudate compounds related to annotated features increased by virocell necromass (benzoic acid, cytidine, dopamine, fructose, glucuronic acid, guanosine, inosine, *p*-coumaric acid, pyridoxine) (**Table S4)**. The compounds were tested at two environmentally/physiologically relevant concentrations (0.5 or 5 mM) on model rhizosphere bacterium *P. putida* infected with its compatible lytic phage gh-1 (MOI 0.01). We compared these metabolite-treated cultures to controls (0 mM compound) and measured changes in optical density (ΔOD_600_) to evaluate compound effects (starting OD_600_ at ∼0.5).

Metabolites that most significantly affected infection dynamics included benzoic acid, guanosine, and, to a lesser extent, dopamine (**Fig. S8**). These were associated with a smaller decrease in ΔOD_600_, indicating less culture clearing by phage lysis compared to infected control cultures without metabolite additions. To control for the effect of phage stock components, we inactivated phage by steam autoclaving and observed the same growth as phage-free cultures. Glucuronic acid, fructose, cytidine, pyridoxine, and inosine did not significantly alter the infection dynamics. Additionally, DMSO used as a solvent for some metabolites did not significantly impact ΔOD_600_ values based on comparisons between +/− DMSO-supplemented cultures (**Fig. S8b**).

To further investigate the effects of benzoic acid and guanosine (the two metabolites with the most significant impact on phage lysis) and *p*-coumaric acid (additional aromatic acid), we supplied these compounds to infected and uninfected cultures of *P. putida*. To evaluate the generality of the metabolite effect on phage infections, we also included a second susceptible soil bacterium, *Bacillus subtilis,* with its lytic phage SPO1. In both cases, phages were added at the MOI of 0.01 to the culture of the host, and ΔOD_600_ was measured (starting OD_600_ ∼0.5). In *P. putida*/gh-1 phage, benzoic acid, guanosine, and *p*-coumaric acid showed reduced culture clearing (**Fig. 4a**), and uninfected *P. putida* had enhanced growth on guanosine and *p*-coumaric acid (**Fig. 4a**). Conversely, *B. subtilis*/SPO1 phage only showed reduced clearing with *p*-coumaric acid, which also inhibited uninfected host growth (**Fig. 4b**). The metabolites did not statistically change phage titer relative to controls as measured by plaque assays (**Fig. S9**).

**Fig. 4:**
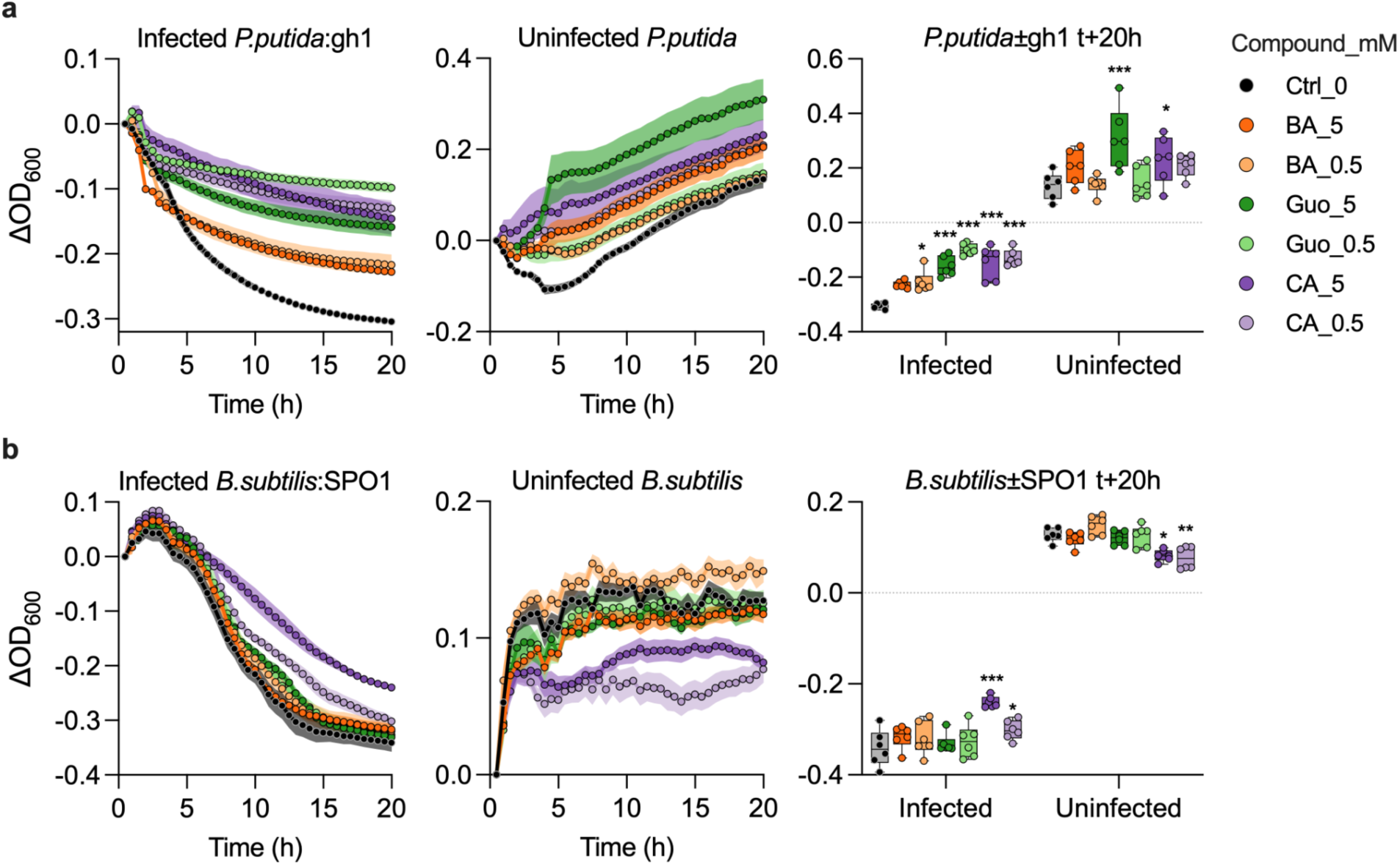
Bioassays showing exometabolites affecting phage infection *in vitro*. Optical density changes ΔOD600 for uninfected or phage-infected cultures (MOI 0.01 in *V* = 200 µl, starting OD600 at ∼0.5) in LB medium with or without metabolites (Ctrl_0) at 0.5 or 5 mM: benzoic acid (BA), guanosine (Guo) and *p*-coumaric acid (CA). Left: infected culture growth curves. Middle: uninfected bacteria growth curves. Right: final data points at t+20h. (**a**) *P. putida* A.3.12 with phage gh-1. (**b**) *B. subtilis* 168 M and phage SPO1. Box plots show all data points, with hinges for the 25th to 75th percentiles, median line, and whiskers for min/max values. Statistical analysis: 2-way ANOVA, *n*=6 (separate mixtures/infections) with the Dunnett test vs. control (ns *p*>0.5, \**p*<0.05, \*\**p*<0.01, \*\*\**p*<0.001).

## 4. Discussion

Here, we compared the effects of virocell vs. sonicated necromass on plant N nutrition. We did not observe a significant impact of necromass on the biomass formation of *B. distachyon*. However, the bacterial necromass caused the production of hundreds of metabolic features in the rhizosphere, with nucleosides showing the most significant increase. Using stable isotope tracing and metabolite analysis, we determined that *B. distachyon* produces guanosine and other metabolites from microbial-derived N. Guanosine and certain organic acids (benzoic and *p*-coumaric acids) from root exudates showed selective effects on phage lysis. This finding suggests a complex interaction between virocells and plant metabolism, highlighting the potential for virocells to influence plant health and nutrient acquisition that are not yet fully understood.

The ^15^N stable isotope tracing experiments showed that there was a small but significant plant uptake of sonicated necromass, while phage lysis contributed no additional N to *B. distachyon* (**Fig. 1c**). This difference can be attributed to qualitative changes in metabolite composition during phage infection (**Fig. S6c**), particularly a reduction in N-containing amino acids and peptides available for plant uptake (Paungfoo-Lonhienne *et al*. 2008; Näsholm *et al*. 2009). These N-containing compounds can theoretically be redirected toward building more phage particles (Jover, Effler, Buchan, Wilhelm & Weitz 2014). Additionally, we calculated that lysis of 3.3e10 *Pseudomonas* cells (≈10 ml of OD 1.4) could theoretically satisfy the N requirements of 4-week-old *Brachypodium* plants (**Table S2**). This exceeds the number of bacteria in the rhizosphere based on a reported 1e9 CFU/g root in 21 days post-inoculation (Lin *et al*. 2023) and the typical bacterial densities in the rhizosphere (Watt, Hugenholtz, White & Vinall 2006). However, these static estimates fail to capture the spatial-temporal dynamic N flow in the rhizosphere, including competition and mutualism between plants and microbes, with rhizodeposits-driven continuous microbial turnover releasing N for plant uptake, complemented by N supply from diazotrophs (Kuzyakov & Xu 2013). Therefore, phage lysis could theoretically contribute to plant N status in the long term, and the importance of this to plant growth might depend on the environment.

Necromass caused a specific increase in rhizosphere metabolites (**Fig. 2b,c**). This finding aligns with the understanding that root exudates are dynamic and can be influenced by various factors, such as plant age, N supply, microbial activity, diurnal cycles, sterility, or sucrose supply, suggesting that necromass could be an additional factor influencing root exudation (Zhalnina *et al*. 2018; Korenblum *et al*. 2020; McLaughlin, Zhalnina, Kosina, Northen & Sasse 2023; Novak *et al*. 2024). The elicited metabolites included the nucleoside guanosine (**Fig. 3**), which can affect virocell growth (**Fig. 4a**). Our stable isotope labeling and metabolomics (**Fig. 3 and 4**) suggest that nucleosides are either produced via rapid recycling of microbial N by plants followed by root exudation or by the activity of plant exogenous nucleases on nucleic acids from necromass. Both processes are possible as previous studies report root exudation of nucleases and guanosine (Chen, Delatorre, Bakker & Abel 2000; Zhalnina *et al*. 2018; McLaughlin *et al*. 2023).

The presence of compounds promoting the growth of bacterial hosts, such as *P. putida,* in the rhizosphere (**Fig. 4a**) is aligned with the observation that the rhizosphere is a hotspot for bacterial growth (Kuzyakov & Blagodatskaya 2015), which subsequently can facilitate replication of phages (Sokol *et al*. 2022). The opposite pattern was observed for *B. subtilis* as a host (**Fig. 4b**), for which compounds providing increased resistance to phage infection were associated with reduced growth of uninfected cells. These contrasted patterns likely hint at different virus-host dynamics, with the former (*P. putida*) associated with the maintenance of an uninfected host pool via fast cellular replication, while the latter (*B. subtilis*) is likely associated with stalled viral infections due to limited host growth. In both cases, however, and especially for *P. putida*, increased bacterial growth despite active phage infections may be beneficial for the plant and a key mechanism to maintain a steady community of beneficial bacterial partners. This plant-driven host growth modulation could explain the reported taxonomic shifts between bulk soil and rhizosphere virome (Bi *et al*. 2021).

This study has several limitations, necessitating additional studies to draw generalizable conclusions. First, the study focuses on bacterial lysates from a single species in a soil-free context, thus omitting the N in insoluble cell debris, soil viral and microbial diversity, and metabolite-mineral associations (Sasse *et al*. 2020; Nayfach *et al*. 2021). Future research should utilize more phage-host combinations, but this is limited by the large fraction of uncharacterized and uncultured soil microbes and their viruses (Rappé & Giovannoni 2003; Mouginot *et al*. 2014; Fierer 2017; Lloyd, Steen, Ladau, Yin & Crosby 2018; Nayfach *et al*. 2021). Sonication can estimate potential total N release without necessitating cultured phage; however, it omits the changes during virocell biochemical remodeling (**Fig. S6**) (Howard-Varona *et al*. 2020). Another complexity in phage inoculum may be unlabeled N (**Fig. 3, Fig. S6**), likely from lysed hosts or phage particles (e.g., capsid and phage DNA), which might influence ^15^N tracing mainly due to non-uniform labeling of the source pool (Stark 2000). The methods of separation of organic compounds from phage inoculum often involve centrifugation and dialysis (Bonilla *et al*. 2016; Luong, Salabarria, Edwards & Roach 2020). However, as shown for T4 phage and *E. coli*, the efficiency of phage-induced lysis is significantly reduced without the metabolic support typically provided by organic compounds (Bryan, El-Shibiny, Hobbs, Porter & Kutter 2016).

This study has several important implications. It provides additional support for the emerging view that while phage activity in the rhizosphere impacts nutrient dynamics, it is unlikely to be a major source of plant N. Importantly, the observed selective elicitation of root exudate metabolites by virocell necromass, several of which impact virocell growth. Future work should investigate the possible benefits of the root exudate metabolites on other phage-bacteria-plant systems and the resulting impact of phage lysis on the biogeochemical cycling of other elements. Namely, the impacts of soil metabolites on the quantitative life history trait data through phage adsorption kinetics and one-step growth experiments could guide predictive modeling efforts (Deng *et al*. 2012). Such investigations are essential for gaining a molecular understanding of soil microbiome functioning, which is crucial for harnessing microbiomes to promote plant growth, sequester C, and improve soil quality (Jansson & Hofmockel 2020; Trivedi, Leach, Tringe, Sa & Singh 2020).

## 5. Material and Methods

### 5.1. Preparation of phage stock and plaque assays

We propagated *Pseudomonas putida* bacteriophage gh-1 (ATCC 12633-B1) using the host *P. putida* A.3.12 (ATCC 12633) (Lee & Boezi 1966). The initial propagation followed the manufacturer-recommended Adams agar-overlay method (Adams 1959), briefly: *P. putida* grew overnight in BD nutrient broth (beef extract 3g/L, peptone 5g/L; BD 234000), then 100 µl of culture was diluted in 2.5 mL of melted 0.5% agar (same medium), which was then poured onto nutrient 1.5% agar plates (BD 213000). After hardening, the plate was evenly covered with 0.5 ml of phage (rehydrated in BD nutrient broth) and incubated at 27°C for two days. Subsequently, the soft agar was scraped, then centrifuged at 12,000 g for 25 minutes, and the supernatants were filtered through a 0.2 µm PVDF Millipore filter. The main plant isotope labeling experiment used the nutrient broth-propagated gh-1 stock (with a 2.1e7 PFU/ml concentration).

To boost phage titer, subsequent phage propagations for bioassays utilized a modified technique by using LB Miller medium (Tryptone 10g/L, Yeast Extract 5g/L, NaCl 10g/L, pH 7.2; Growcells MBLE-7030) instead of BD nutrient broth and by mixing 100 µl of host and phage stock directly into melted overlay agar which was then incubated overnight. Phages were harvested by disrupting the overlay agar with 6 ml of SM buffer (50 mM Tris-HCl, 8 mM magnesium sulfate, 100 mM sodium chloride, 0.01% gelatin, pH 7.5; Thermo Scientific, J61367-AK), followed by centrifugation at 8000 g for 10 min and filtration of supernatant via 0.2 µm PES syringe filters. The modified technique was used to propagate the gh-1 phage with host *P. putida* A.3.12 and SPO1 phage (ATCC 27370-B1) with host *B. subtilis* 168M (ATCC 27370), yielding routinely 3e9 PFU/mL for gh-1 and 5e7 PFU/mL for SPO1. All phage were stored as filtered lysates at 4°C. Phage titer (PFU/ml) was estimated by spotting a 10-fold serial dilution of each phage in SM buffer supplemented with 10 mM CaCl_2_ on a lawn of a compatible host using the top agar overlay plaque assays with LB 0.5% agar.

### 5.2. Preparation of bacterial necromass media

A culture of *Pseudomonas putida* A.3.12 (ATCC 12633) grew on M9-Celtone medium containing 42.3 mM Na_2_HPO_4_, 22.0 mM KH_2_PO_4_, 8.6 mM NaCl, 15 mM glucose, 2 mM MgSO_4_, and 0.1 mM CaCl_2_. In addition, the unlabelled medium contained 1g/L of NH_4_Cl and 1g/L Celtone® Base Powder (Cambridge Isotope Laboratories, Inc., product number CGM-1030P-U). The ^15^N-labeled medium contained 1g/L ^15^NH_4_Cl (≥98 atom % ^15^N) and 1g/L of Celtone® Base Powder (≥98% ^15^N) (Cambridge Isotope Laboratories, CGM-1030P-N). Both media pH were 7. A 50 ml culture of the bacteria was incubated for 24 h at 30°C in 250 ml Erlenmeyer flasks under constant shaking at 200 rpm, reaching OD_600_ of 1.515 in unlabelled and 1.537 in the ^15^N-labeled cultures.

The extracellular inorganic N and exometabolites were removed by centrifuging 30 ml of cultures for 10min/4000g/RT. Then we discarded the supernatant, washed the pellet with 30 ml of half-strength Murashige and Skoog basal salts medium without nitrates (N-free ½ MS salts) (Caisson lab, MSP10), re-pelleted for 10min/4000g/RT, discarded supernatant, and finally resuspended in 30 ml of N-free ½ MS salts. We split the washed cells equally into 3 tubes. The first tube containing 10 ml of cell suspension was inoculated with 200 µl of gh-1 phage stock (4.2×10^6^ PFU / 3.77×10^10^ CFU = MOI 0.0001) and incubated at 30°C under constant shaking at 200 rpm for 1h to ensure at least two infection cycles (21 min latency period) (Lee & Boezi 1966). We sonicated the second tube with 10 ml of cell suspension for 5 min on ice using a probe sonicator (Q125, Qsonica L.L.C, Newtown, CT, USA) set to 60% amplitude and 50s pulse and 10s break interval. The third 10 ml aliquot of unlysed cells stayed at 30°C for 1h. We saved 5 µl aliquots of each of the 3 treatments for cell density determination. To prevent clotting during filtration, we centrifuged the cell suspensions for 5 min/1000g/RT and collected the supernatant. We adjusted the supernatants to 100 mL (final 0.1x concentration) using an N-free ½ MS salts solution. This 0.1x concentration resulted in sonicated lysed cells equivalent to 3.3e8 cells/mL (≈ OD_600_ 0.1). This concentration approximates the average bacterial cell densities at 1e8 CFU/g of a potting substrate in the rhizosphere of maize or *Brachypodium* (Di Battista-Leboeuf, Benizri, Corbel, Piutti & Guckert 2003; Lin *et al*. 2023). Then, we filtered the suspensions using a 0.2 µm PES vacuum filtration system. We used sterile N-free ½ MS salts medium filtered via 0.2 µm PES membrane for plant axenic and technical controls. Half of each filtrate resulted in 5 replicates for analysis at t_0_. The second half of each filtrate yielded 5 replicates for plant assays. We prepared a gh-1 baseline control for t_0_ analyses by adjusting 100 µl of gh-1 phage stock to 50 ml with N-free ½ MS salts and filtering the solution via 0.2 µm PES membrane. We used 7 ml of the t_0_ media for LC-MS and 1 ml for TOC.

### 5.3. Cell density enumeration and calibration curve

We determined cell density (cells/ml) using a disposable hemocytometer (C-chip DHC-S02-2, Incyto) combined with phase-contrast microscopy under 400x magnification. First, we 10x diluted an aliquot of cells lysed by phage (virocell), sonicated, and unlysed to 50 µl with N-free ½ MS salts; then, we counted the number of cells in the hemocytometer chamber equal to the volume of 200×10^-7^ cm^3^. The calculation for cell concentration was cells per ml = no of cells × dilution factor × volume factor. The volume factor = 1/(200 × 10^-7^ cm^3^) = 50,000. To convert between OD_600_ and the cell count of *P. putida* A.3.12, we created a linear correlation with an intercept set to 0. Dilutions of the culture were measured for OD_600_, and cell count was quantified through microscopy and a hemocytometer.

### 5.4. Plant growth, harvest, and medium sampling

John Vogel (JGI, CA, USA) provided seeds of *Brachypodium distachyon* Bd21-3. We surface-sterilized the seed with 70% ethanol for 30 seconds, then with 6% w/v sodium hypochlorite solution for 5 min, and washed the seeds 5x with sterile milli-Q water. Seeds were then plated on plates containing 1.5% phytoagar in ½ MS basal salts and put into the dark at 4°C for 3 days. Seeds on plates were then germinated vertically in a growth chamber (CU36L4, Percival Scientific Inc, Perry, IA, USA) set to PPFD at 150 µmol/m^2^/s, 12 h dark/light cycle, and continuous 22°C. These settings were used for all subsequent plant growth experiments. After three days, we transferred the seedlings to fabricated ecosystems (EcoFAB 2.0, https://eco-fab.org/). EcoFAB 2.0 is a plant growth system that allows reproducible rhizosphere studies of plant traits and exometabolites (Novak *et al*. 2024). The EcoFABs 2.0 units were assembled, sterilized, and aseptically filled with 10 ml of ½ MS salts (MSPO1, Caisson lab) at a pH of 5.74, followed by seedlings transfer (Andeer & Novak 2023; Novak *et al*. 2024).

We placed the EcoFAB 2.0 devices into a growth chamber set to the germination settings. The medium was replaced weekly with fresh ½ MS salts solution. After 3 weeks of growth, we removed spent medium and supplied plants with 10 ml of ^15^N-labeled or unlabeled filtered media consisting of i) virocell (2.3e8 lysed cells ml^-1^), ii) sonicated cells (3.3e8 lysed cells ml^-1^), iii) unlysed cells, or iv) cell-free ½ MS medium (axenic plant treatment). In addition, we included treatment v) technical control that was EcoFAB 2.0 filled with cell-free ½ MS salts medium as the baseline for metabolomics. Each treatment had 5 biological replicates. After one week (t_1_), we collected the spent media, plant shoots, and roots. We adjusted the spent media to 10 ml, filtered it via 0.2 µm PES membrane syringe filter (4652, Pall Corporation), and used 7 ml for metabolomics and 1 ml for TOC. To check for sterility, we plated 10 µl of the plant spent media from the end of weeks 3 and 5 on LB agar plates incubated at 27°C for 7 days. No bacterial colonies grew on the plates, indicating 100% sterility. The root and shoot tissues were frozen at −80 °C and then lyophilized (Labconoco FreeZone 2.5) to determine dry weight.

### 5.5. Plant-dependent necromass turnover experiment

A follow-up unlabeled necromass turnover experiment was performed to check for the formation of metabolites in 0.1x dilution of sonicated *P. putida* lysates without plants in EcoFAB 2.0 after one week of incubation (t_1_). The controls in EcoFAB 2.0 were 0.1x lysate at t_0_, *B. distachyon* supplied with 0.1x lysates at t_1_, and N-free ½ MS salts technical control at t_1_. The setup, timing, growth conditions, and medium sampling followed the experiment above, with each treatment having *n*=5 biological replicates.

### 5.6. Quantification of total N and C in necromass media

To quantify the concentration of N and C in the necromass media, the 1 ml aliquots of t_0_ and t_1_ medium samples were adjusted to 9 ml with milli-Q water and analyzed for total nitrogen (TN) and dissolved organic carbon (DOC) using a TOC analyzer (ASI-L autosampler coupled to TOC-L H-type with TNM-L, Shimadzu, Kyoto, Japan). The DOC was measured as non-purgeable organic C. Samples were acidified at 1.5% v/v with 1M HCl to convert inorganic C to carbonates and then sparged with ultra-high purity synthetic air for 2.5 minutes at 80 mL/min to remove volatile organic C and carbonates, then 3–5 injections of 50 µL (within 2% CV) were measured for C and N with a maximum integration time of 4:50 min. One needle wash was performed for each sample. Standard curve and calibration check solution were prepared with potassium hydrogen phthalate and potassium nitrate. Millipore Milli-Q lab water was used to blank the system.

### 5.7. Stable isotope mass spectrometry and uptake of microbial-N into plant tissue

The dry root and shoot tissues were homogenized using two metal beads and a beat mill (MM 400, Retsch GmbH, Haan, Germany) set to 30 Hz for 10 min. The dried powdered tissue (1-3 mg) was then weighed into tin capsules 8 x 5mm (D1008, EA Consumables) and measured for N elemental (% dry weight) and stable isotope composition (^15^N at-%) at The Stable Isotope Facility, University of California, Davis. The analytical system consisted of an Elementar vario MICRO cube elemental analyzer (Elementar Analysensysteme GmbH, Langenselbold, Germany) interfaced with a Sercon Europa 20-20 IRMS (Sercon Ltd., Cheshire, United Kingdom). Raw isotopic values were adjusted for changes in linearity and instrumental drift and normalized using in-house reference materials calibrated against international reference materials traceable to the primary isotopic reference material, i.e., Air. Elemental totals were calculated based on IRMS peak area using linearity references as a calibration curve. Precision (σ) and accuracy for ^15^N (at-%) measured on nylon powder were <0.003 % and <0.001 %, respectively. The analytical range was 20–150 μg N, and the limit of quantification (LOQ), based on the peak area, was 20 μg N. The uptake of microbial-N into plant root or shoot tissues was calculated according to equation 1 (Stark 2000):

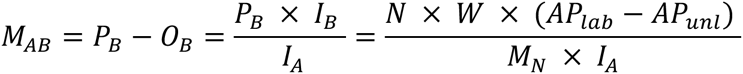

where *M*_AB_ is the total amount of N that flowed from labeled microbial source pool A to plant sink pool B. *P*_B_ is the amount (in mol) of N in the plant sink pool (B). *O*_B_ is the other-N (non-labeled) in the plant N pool. *I*_B_ is at-% excess of plant pool B. *I*_A_ is at-% excess of source pool A, assuming the enrichment of pool A at 98% ^15^N and the natural abundance of ^15^N at 0.3663% (Fry 2006), then the *I*_A_ = 97.6337%. *N* is N content in plant tissue (wt %). *W* is plant dry tissue weight. *AP*_lab_ is the ^15^N at-% of labeled plant tissue, while *AP*_unl_ is the ^15^N at-% of the unlabeled plant tissue. *M*_N_ is the molar weight of N at 14.0067 g mol^-1^.

### 5.8. LC-MS/MS sample preparation and analysis

Samples were lyophilized until dry and stored at −80°C until extraction. Dried samples were extracted in 1 mL of LC-MS grade methanol chilled at −20 °C, vortexed twice for 10 s, and sonicated in a water bath sonicator with ice for 15 min. Samples were centrifuged at 10,000g/5 min/10°C to pellet salts. The supernatant was collected, transferred to new microcentrifuge tubes, and dried using a SpeedVac vacuum concentrator (Thermo Scientific Savant SpeedVac SPD130DLX) at RT. Dried samples were stored at −80°C until final resuspension. Samples were resuspended in 150 µL −20°C chilled methanol containing internal standards (**Table S1**) and were vortexed, sonicated, and centrifuged using the same settings mentioned above. The supernatant was transferred to Ultrafree MC-GV centrifugal filters with Durapore PVDF 0.22 µm pore size filters (Millipore) and centrifuged at 10,000g/5min/10°C. The filtrate was then transferred to amber glass vials with inserts for LC-MS/MS analysis. Metabolites from samples were separated using hydrophilic liquid interaction chromatography and detected using a Thermo QExactive Hybrid Quadrupole-Orbitrap Mass Spectrometer. Data-dependent MS2 fragmentation spectra were collected from the top two most intense ions not previously selected for fragmentation within 7 s. LC-MS/MS parameters are available in **Table S1**.

### 5.9. Metabolomics data analysis

The data from the labeling experiment were analyzed by untargeted metabolomics followed by spectral matching to an online MS2 database. MZmine 2.0 (**File S1 and S2**) generated a list of features (Pluskal, Castillo, Villar-Briones & Oresic 2010), resulting in 3708 and 4572 features for negative and positive modes, respectively. When available, each feature’s most intense fragmentation spectrum was uploaded to the GNPS (Global Natural Products Social Molecular Networking) (Wang *et al*. 2016). A match between a feature and compound spectrum (GNPS settings in **Table S1**) in the GNPS database resulted in putative annotations. Downstream filtering accepted only features with RT>0.6 min, maximum peak height > 1e6, and a maximum sample peak intensity >10x the maximum in extraction and technical controls. These filtered features (2059 after merging polarities, **Table S3**) were used for untargeted data analysis. NPClassifier was used to determine metabolite classification (Kim *et al*. 2021). The results for isotopically unlabeled samples were used to create figures. To identify the number of intersecting features between treatments, we applied a peak intensity threshold of >1e6.

For the necromass turnover experiment, metabolites were identified by searching the data using an in-house library of m/z, RT, and MS2 fragmentation data from authentic reference standards using Metabolite Atlas (https://github.com/biorack/metatlas) (Sumner *et al*. 2007; Bowen & Northen 2010). We included metabolites with a maximum sample peak height >10x of extraction and technical controls. The identified metabolites were manually classified using the PubChem Classification Browser (Kim 2021).

The data from the labeling experiment were used for targeted metabolomics to determine ^15^N labeling levels in N-containing metabolites. Metabolites were identified using the Metabolite Atlas, and peak height values for all theoretical ^15^N isotopologues were extracted. Isotopologues with significant confounding background signals were excluded before quantification. M0 intensities, representing the ^14^N monoisotopic m/z, were calculated as the difference between total vs labeled intensity (M1+n). The M1+n intensities were the sum of all other isotopologue peak heights. For downstream processing, we selected ^15^N-supplied treatments and filtered metabolites with a maximum total intensity >1e6 and average percent labeling >50% in at least one treatment group.

### 5.10. Quantification of N forms in sonicated bacterial lysates

To evaluate the release of N from lysed bacterial cells for plant uptake, we quantified nitrate (NO_3_-N) and ammonium (NH_4_-N) alongside protein (protein-N) in sonicated cell lysates and their spent media. We selected soil and root-colonizing bacteria *Pseudomonas putida* KT2440 (ATCC 47054), *Pseudomonas simiae* WCS417 (obtained from Benjamin Cole, Joint Genome Institute), and *Pseudomonas putida* A.3.12 (ATCC 12633) (Molina 2000; Cole *et al*. 2017; Liffourrena & Lucchesi 2018). The bacteria grew on LB agar plates at 27°C in the dark for 1 day. Then, we aseptically resuspended a single colony in 50 µl of 1x PBS (AM9624, Invitrogen) and inoculated 1 µl of the cell suspension into 30 ml of unlabeled M9-Celtone medium in Erlenmeyer flasks (*n*=3 biological replicates). The culture grew for 24 hours at 27°C at 200 rpm; then, the growth was measured as OD_600_. We collected 10 ml of each culture to sample the spent medium and create lysates. The spent culture medium was collected by centrifugation at 4000g/10min/RT and filtered via 0.2 µm PES membrane syringe filter (4652, Pall Corporation). The pellets were washed with 1x PBS, re-pelleted at 4000g/10min/RT, and resuspended in 10 ml of Milli-Q water. The cell suspension was sonicated on ice for 5 min with a probe sonicator (Q125, Qsonica L.L.C, Newtown, CT, USA) set to 60% amplitude and 50s pulse and 10s break interval, and the OD_600_ was measured to estimate the lysis efficiency. Centrifugation of the lysates at 4000g/5min/4°C removed cell debris, and then the supernatant was filtered via a 0.2 µm PES membrane syringe filter (4652, Pall Corporation). The NH_4_-N was quantified using two photometric Ammonium Tests (Supelco 100683 Merck, range 2.0–150.0 mg/L NH₄-N for medium samples and Supelco 114752 Merck, range 0.01–3.00 mg/l NH₄-N for lysate samples) at 690 nm against a linear calibration curve of ammonium standard (59755, Sigma-Aldrich). The NO_3_-N was quantified using a photometric Nitrate Test (Supelco 109713 Merck, range 0.1–25.0 mg/L NO_3_-N) at 340 nm against a linear calibration curve of nitrate standard (74246, Sigma-Aldrich). For protein quantification, we used a photometric Pierce™ BCA Protein Assay Kit (23227, Thermo Scientific, range 20–2000 µg/mL) at 562 nm against a linear calibration curve of kit-included Albumin (BSA) standard. To convert protein concentration to protein-N concentration, we applied a standard approximation of 16% N content in protein (Maclean *et al*. 2003). Calculated concentrations that were negative were assumed to be 0.

### 5.11. *P. putida* A.3.12 influence on *B. distachyon* growth

As indicated above, the EcoFAB 2.0 with *B. ditachyon* was set up with ½ MS salts. Then, 4 days after seedling transfer, *P. putida* A.3.12 (pre-washed and resuspended in ½ MS salts) was inoculated to an OD600 of 0.01. The controls were uninoculated plants. Each group had *n* = 5 biological replicates. Weekly water additions were made to offset evaporation. After 22 days post-inoculation (DAI), plant growth was measured by weighing fresh shoots and root biomass. The sterility of the EcoFAB 2.0 was checked by incubating 2 µl of growth medium on LB agar plates for 7 days.

### 5.12. Long-term effect of necromass on *B. distachyon* growth

To study the long-term effect of necromass on plant development, we grew *B. distachyon* in EcoFAB 2.0 for 2 weeks on ½ MS salts medium, which was then replaced for the treatment groups with sonicated unlabeled *P. putida* necromass at 0.1x or 0.2x concentrations (resuspended in N-free ½ MS) for an additional 3 weeks. Control treatments included plants with N-free ½ MS (-N) or ½ MS salts (+N). Each group had *n* = 5 biological replicates. The evaporated water was re-supplied weekly. Plant biomass was lyophilized to measure root and shoot dry weight.

### 5.13. Effect of exometabolites on phage lysis *in vitro*

Two initial bioassays investigated the effects of selected root exudate metabolites on phage lysis using the gh1 phage with the host rhizosphere-dwelling bacterium *P. putida* A.3.12 (Lee & Boezi 1966; Liffourrena & Lucchesi 2018). Two initial bioassays included root exudate metabolites dopamine, pyridoxine, benzoic acid, cytidine, fructose, glucuronic acid, guanosine, and inosine (**Table S4**) (Lee, Lee & Lee 2006; Liu *et al*. 2015; Kawasaki *et al*. 2016; Zhalnina *et al*. 2018; Baker *et al*. 2022; McLaughlin *et al*. 2023; Novak *et al*. 2024). The two follow-up bioassays tested benzoic acid, another aromatic carboxylic acid from root exudates—*p*-coumaric acid (Novak *et al*. 2024), and guanosine with either uninfected or phage-infected host of *P. putida* A.3.12 with gh1 phage or another rhizosphere-dwelling bacterium *B. subtilis* 168M (ATCC 27370) with lytic phage SPO1 (ATCC 27370-B1) (Okubo *et al*. 1964; Gallegos-Monterrosa *et al*. 2016). The bacterial pre-cultures grew in Miller’s LB broth (Growcells) overnight at 27°C. Then, the spent medium was removed, and the cells were resuspended in fresh Miller’s LB medium. Then, cultures were loaded onto 96-well plates (Costar), and phages, solvents, and metabolites were mixed in each well. We used MOI at 0.01, which involved adding 1 µl of gh-1 inoculum (3e9 PFU/ml) or 57 µl of SPO1 inoculum (5e7 PFU/ml) to 2.4e8 host cells in a final volume of 200 µl, which equals to starting OD_600_ at ∼0.5 and phage titer at ∼1.5e7 PFU/ml. The metabolites were added at 0.5 and 5 mM to reflect physiologically relevant levels for bacterial growth (Novak *et al*. 2024) and to cover the upper range of soil metabolite concentrations, which can vary from 0.03– 70 µM in bulk soil (1:1 water extract) to 2.7–12.9 mM in hotspots (for amino acids) (Jenkins *et al*. 2017; Hill, Jones, Paterson & Hill 2019). Benzoic acid, guanosine, and *p*-coumaric acid were dissolved in 5% dimethyl sulfoxide (DMSO) (NWR Life Sciences), while the other compounds were water-soluble. Our experiments included multiple controls without metabolite (0 mM) in the first bioassay: infected bacteria, uninfected bacteria, bacteria with inactivated phage (steam autoclaved), and uninoculated LB medium. The second bioassay had the same controls, with or without 5% DMSO. The two follow-up experiments had controls of infected and uninfected bacteria with 0 mM metabolite and 5% DMSO. To control for the SM buffer in the phage treatments, the controls without phage received an equal volume of SM buffer. Each treatment had *n*=6 biological replicates (separate mixtures/infections). The plates were covered with a Breath-Easy® sealing membrane (Sigma-Aldrich) and incubated at 27°C in the Synergy H1 microplate reader (Biotek). The OD_600_ of the plates was measured every 30 min for 20 h. Then, the ΔOD_600_ was calculated as a difference between OD_600_ at the start and subsequent time points. Afterward, we selected 3 random replicates from infected treatments to compare phage titers by plaque assays, focusing on benzoic acid and guanosine for *P. putida* and p-coumaric acid and guanosine for *B. subtilis*.

### 5.14. Statistical analyses and software

RStudio version 1.4.1106 generated the upset plots and Venn diagrams to visualize intersecting sets using the UpSetR and ggVennDiagram packages (Core 2016; Conway, Lex & Gehlenborg 2017; Gao, Yu & Cai 2021). The heat maps were generated with RStudio version 4.0.5 using heatmap.2 in the gplots package (Warnes *et al*. 2024). GraphPad Prism 10 version 10.2.2 was used to generate all other plots and statistical analyses. We calculated the fold-change and *p*-value for volcano plots in Microsoft Excel version 16.59. We calculated the fold change (fc) as fc = (average peak height at t_1_ + 1)/(average peak height at t_0_ + 1). A t-test between the t_0_ and t_1_ time points (*n*=5) determined the *p*-value. The experimental design figure was created using Biorender.com.

## 6. Acknowledgment

VN, SMK, KZ, and TRN gratefully acknowledge funding from the U.S. Department of Energy’s Office of Science, Office of Biological and Environmental Research grant to UC San Diego (DE-SC0021234) and subcontract to Lawrence Berkeley National Laboratory (contract No. DE-AC02-05CH11231). TH, KBL, and SR are supported by the U.S. Department of Energy Joint Genome Institute, a DOE Office of Science User Facility supported by the Office of Science of the U.S. Department of Energy operated under Contract No. DE-AC02-05CH11231. MBS, RN, and VKM were supported by the U.S. Department of Energy’s Office of Science, Office of Biological and Environmental Research as part of a project led by Ohio State University (grant# DOE-BER-248445). Data analysis utilized resources from the National Energy Research Scientific Computing Center (contract DE-AC02-05CH11231).

## 7. Authors contribution

VN conceptualized the study, conducted experiments, collected and visualized data, interpreted findings, and wrote the original draft. MVW conducted experiments and collected data. RN conceptualized the study and provided materials. SMK validated data and reviewed the original draft. TH and KL analyzed the data and reviewed the original draft. SJG and SR interpreted data and reviewed the original draft. KZ and MBS interpreted data, provided funding, and reviewed the original draft. VKM reviewed the methodology, provided materials, interpreted data, and reviewed the original draft. TRN conceptualized the study, interpreted data, provided funding, and reviewed the original draft. All authors edited and approved the final version.

## 9. Data availability

GNPS-negative and -positive modes for the polar metabolite analysis (HILIC) are available at https://gnps.ucsd.edu/ProteoSAFe/status.jsp?task=bcf54efe594f4240acc7e7a919130a33 and https://gnps.ucsd.edu/ProteoSAFe/status.jsp?task=0754b77074864415b55d3c6f6483d332. The LC-MS files are available as a MassIVE dataset under ID number MSV000095290 at https://massive.ucsd.edu/ or via https://doi.org/10.25345/C5QZ22V3X. All raw data and metabolite identification tables are available at https://doi.org/10.6084/m9.figshare.26299084.

