## Supplementary material for "Insights into the role of root exudates in bacteriophage infection dynamics": Figures_S1-S9

**Affiliations:** [1] Environmental Genomics and Systems Biology, Lawrence Berkeley National Laboratory, Berkeley, CA 94720, USA. [2] The DOE Joint Genome Institute, Lawrence Berkeley National Laboratory, Berkeley, CA 94720, USA. [3] Center of Microbiome Science, Ohio State University, Columbus, OH, USA. [4] Department of Microbiology, Ohio State University, Columbus, OH, USA. [5] Department of Civil, Environmental and Geodetic Engineering, Ohio State University, Columbus, OH, USA. [6] EMERGE Biology Integration Institute, Ohio State University, Columbus, OH, USA. [7] Department of Pediatrics, School of Medicine, University of California San Diego, La Jolla, CA 92093, USA. [8] Department of Bioengineering, University of California San Diego, La Jolla, CA 92093, USA. [9] Center for Microbiome Innovation, University of California, San Diego, La Jolla, CA 92093, USA. [10] Program in Materials Science and Engineering, University of California, San Diego, La Jolla, CA 92093, USA.

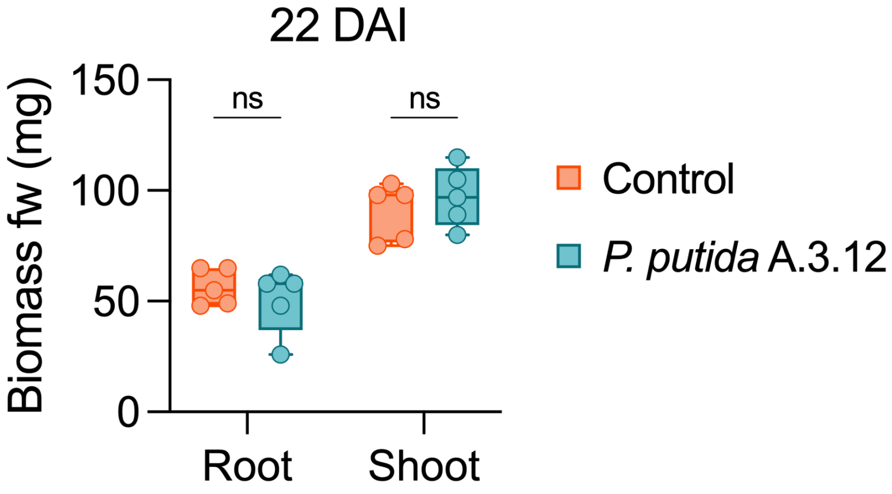


**Fig. S1: *B. distachyon* Bd21-3 interaction with *Pseudomonas putida* A.3.12.** Root and shoot fresh weight of inoculated plants compared to sterile control plants, showing no growth inhibition by the bacteria in 22 days after inoculation (DAI). Statistical analysis was conducted using two-way ANOVA with Sidak's test; ns indicates $p > 0.05$ ($n$=5).

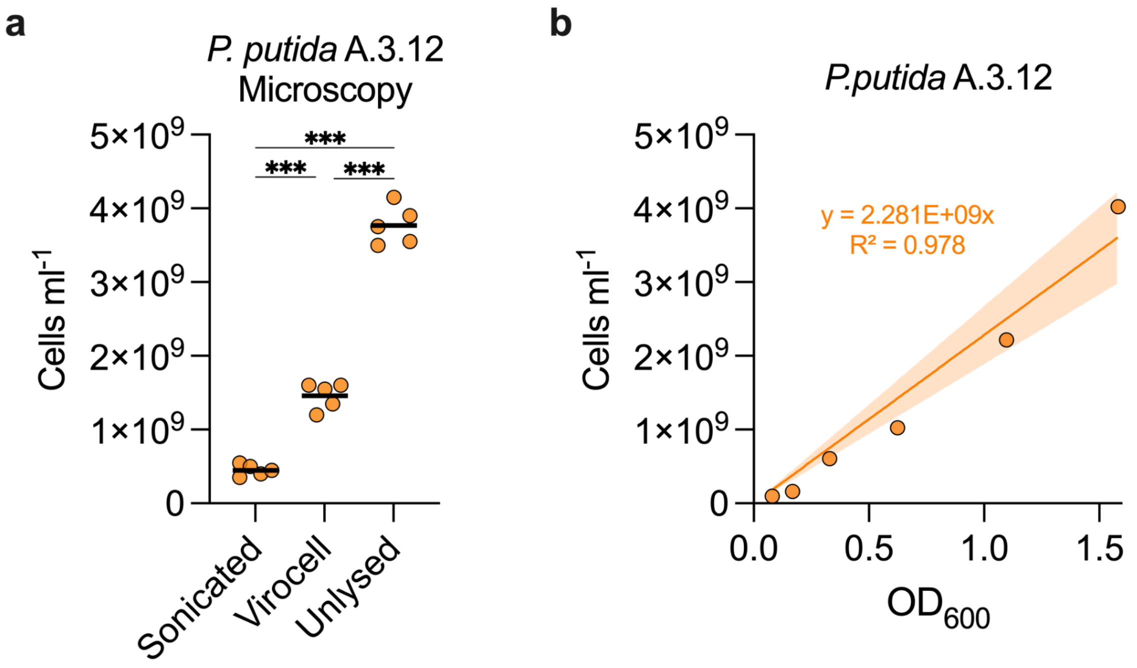


**Fig. S2: Enumeration of cell densities in *P. putida* A.3.12 cultures.** (**a**) Microscopy analysis of unlabeled undiluted cultures showed reduced cell numbers in sonicated and virocell treatments compared to untreated cells, $n$ = 5 technical replicates. One-way ANOVA ($n$=5) with Tukey's test, $n$=5, ***$p$<0.001. (**b**) Linear correlation between cell numbers enumerated by microscopy and optical density at 600 nm ($OD_{600}$).

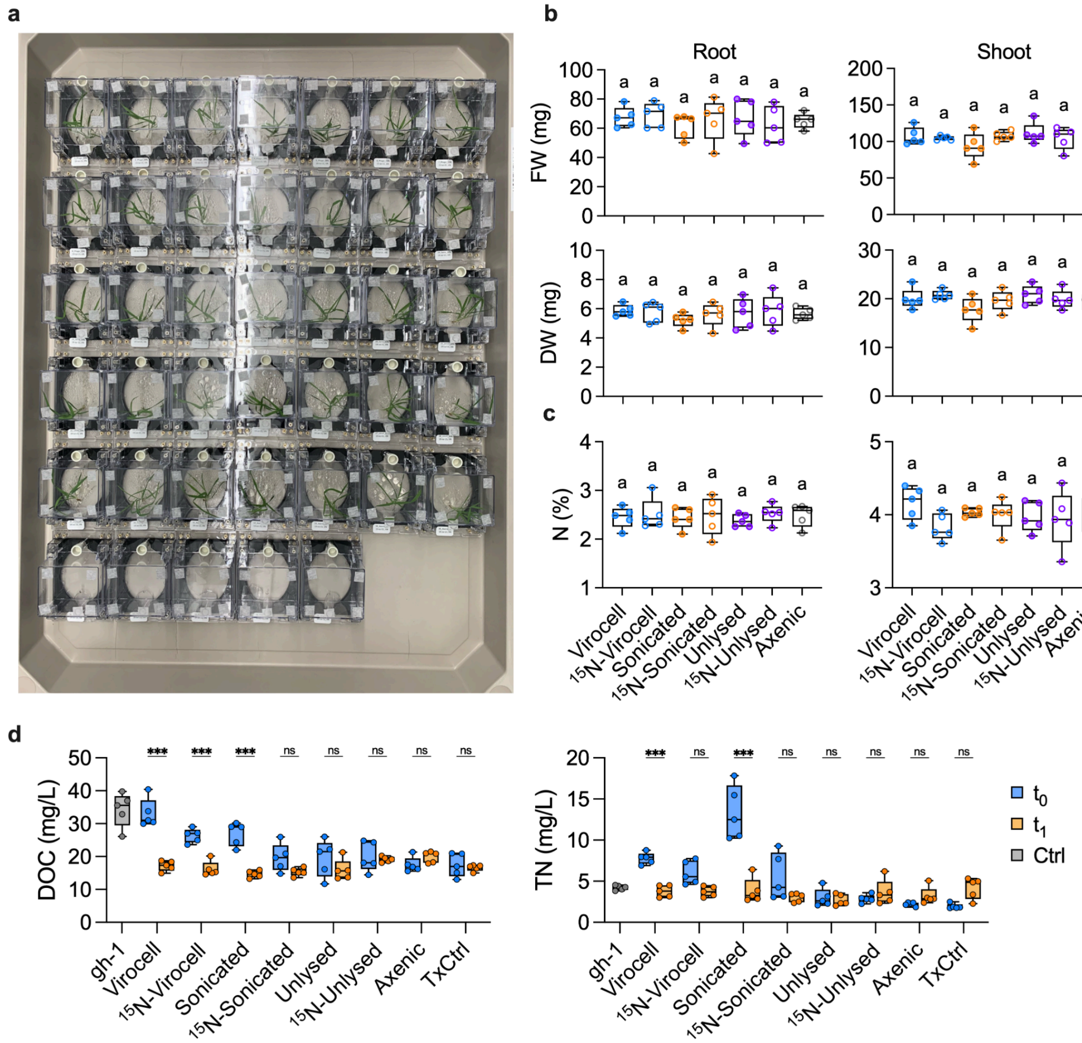


**Fig. S3: Plants phenotype and medium analysis from stable isotope labeling experiment**. (**a**) Photo shows all treatments of *Brachypodium distachyon* plants in EcoFABs 2.0 at the harvest time ($t_1$); each EcoFAB is 127.8 mm L x 85.5 mm W. There was no difference between treatments in (**b**) root and shoot fresh or dry weights or (**c**) total N content in dry tissue (w/w) after one week. Different letters indicate significant differences at $p$<0.05, One-way ANOVA with Tukey's test, $n$=5. (**d**) Total N (TN) and Dissolved Organic Carbon (DOC) in the medium before (t0, blue) and after one week (t1, orange) incubation with plants relative to technical control (TxCrl) and phage gh-1 control (gray). Statistical analysis employed 2-way ANOVA with Šídák test, $n$=5 (*$p$<0.05, **$p$<0.01, ***$p$<0.001). Box plots display all data points, with hinges spanning the 25th to 75th percentiles, a central line denoting the median, and whiskers reaching the minimum and maximum values.

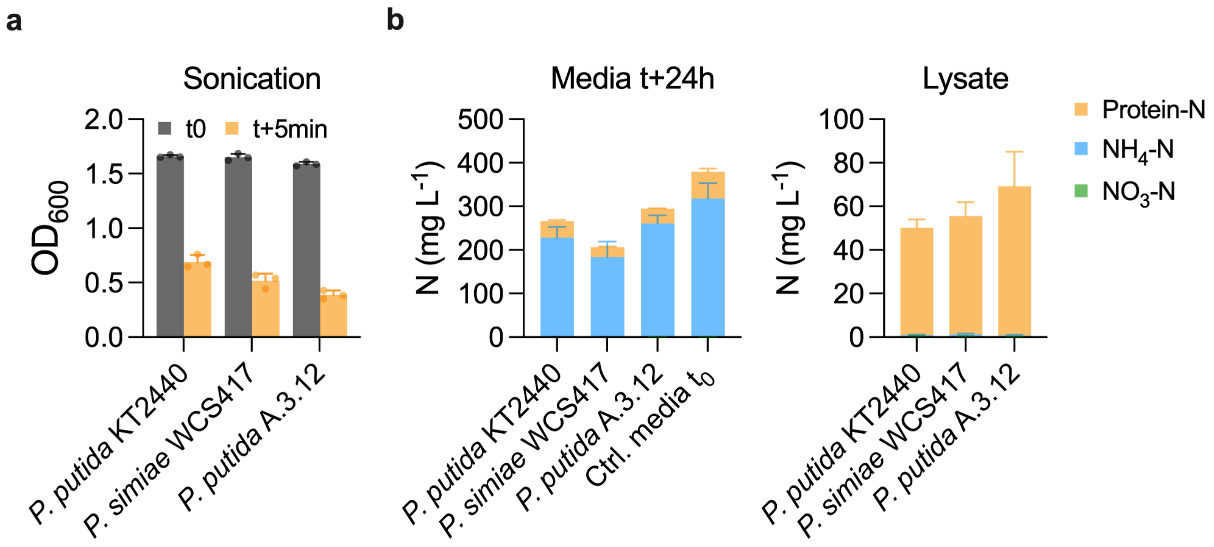


**Fig. S4**: **Nitrogen release from sonicated cells of *Pseudomonas***. (**a**) Optical density $OD_{600}$ for cultures before (t0) and after sonication (t+5min). (**b**) $NH_4$-N, $NO_3$-N, and Protein-N content in spent culture media (from supernatant after centrifugation) after 24h of bacterial cultivation (left) and in filtered bacterial lysates (from sonicated cell pellets) (right). All bar plots show mean with SD, $n$=3 biological replicates.

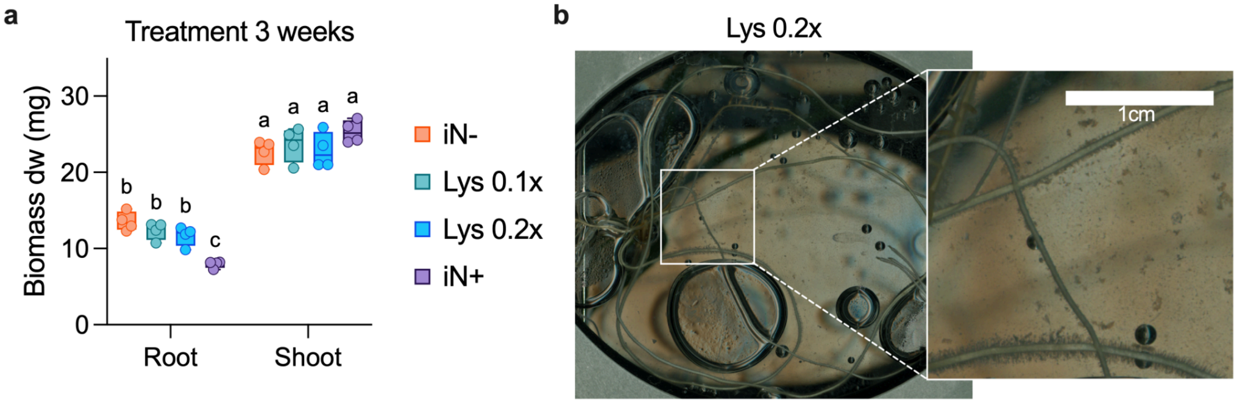


**Fig. S5: Effect of time and dose on plant biomass supplied with sonicated necromass.** Fresh sonicated and filtered lysate of *P. putida* A.3.12 was supplied to *B. distachyon* plants. (**a**) Dry-weight biomass of plants grown with lysates at two dilutions 0.1x or 0.2x for 3 weeks. N-deficient (iN-) and N-containing (iN+) ½ MS basal salts medium were used as controls. (**b**) The picture shows precipitate that formed in the rhizosphere of 0.2x lysate-supplied plants in EcoFAB 2.0. Box plots display all data points, with hinges spanning the 25th to 75th percentiles, a central line denoting the median, and whiskers reaching the minimum and maximum values. Different letters indicate significant differences at $p<0.05$, Two-way ANOVA with Tukey's test, $n=4$.

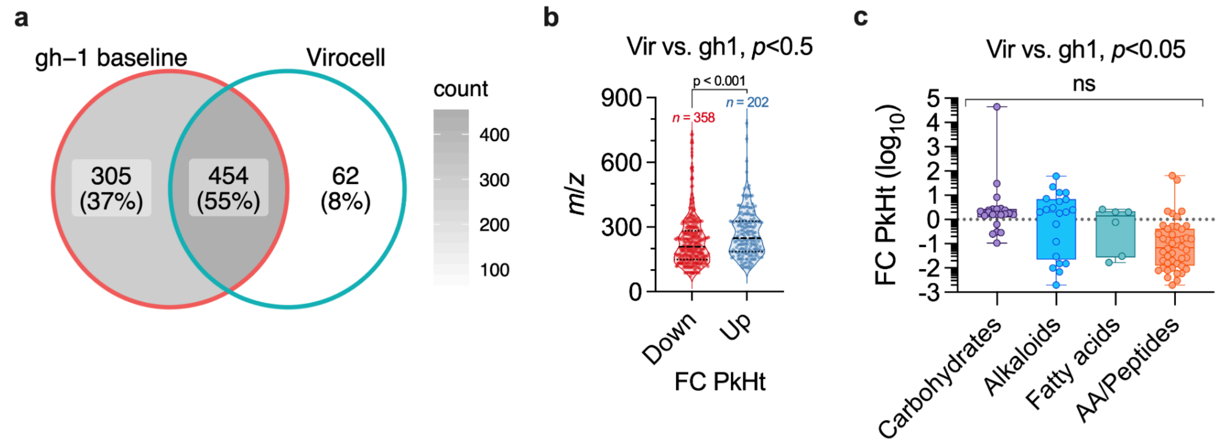


**Fig. S6**: **Exometabolite changes caused by phage gh-1 infection in *P. putida* A.3.12 culture.** (**a**) The Venn diagram shows the number of shared features between virocell and gh-1 baseline (solution of phage inoculum) after 1 hour of infection. (**b**) The *m*/*z* of features significantly higher (up) or lower (down) between virocell and gh-1 phage (t-test, *n*=5). (**c**) Fold change (FC) peak height (PkHt) of annotated features (MQScore>0.7) between virocell and gh-1 phage grouped by biosynthetic pathway (determined by NpClassifier).

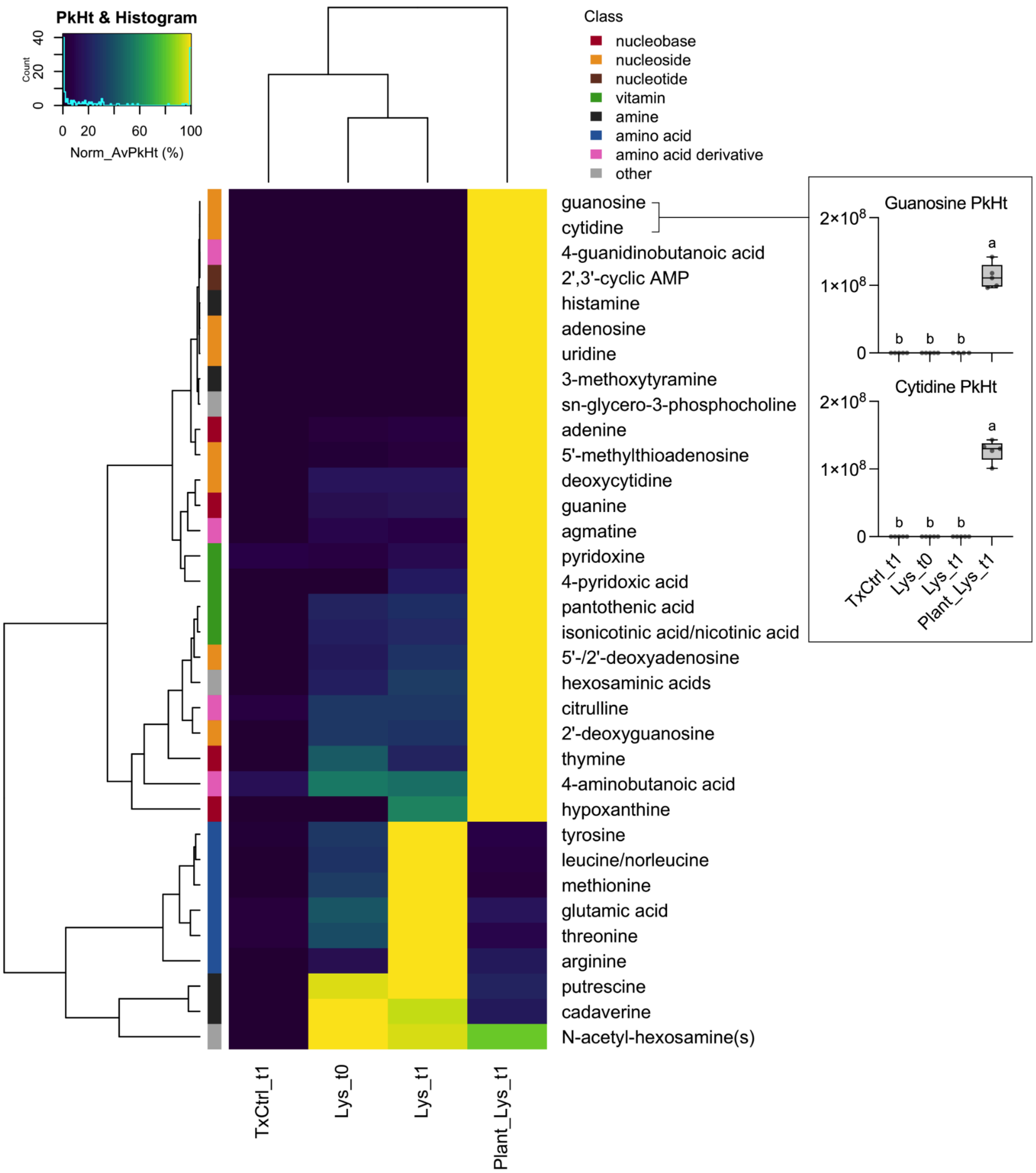


**Fig. S7: Metabolite formation from necromass with and without plants.** Heatmap shows polar metabolites (HILIC-pos) peak height between bacterial lysates incubated with (Plant_Lys_t1) and without plant (Lys_t1) for one week and compared to starting lysate (Lys_t0) and medium-only control (TxCtrl_t1). Amino acid clustering occurred in lysates without plants, while a larger diverse metabolite cluster formed with plants. Box plots highlight plant-driven increases in guanosine and cytidine, displaying all data points, 25th to 75th percentiles, median, and min–max values. Different letters indicate significant differences at $p<0.05$, One-way ANOVA with Tukey's test, $n=5$ biological replicates.

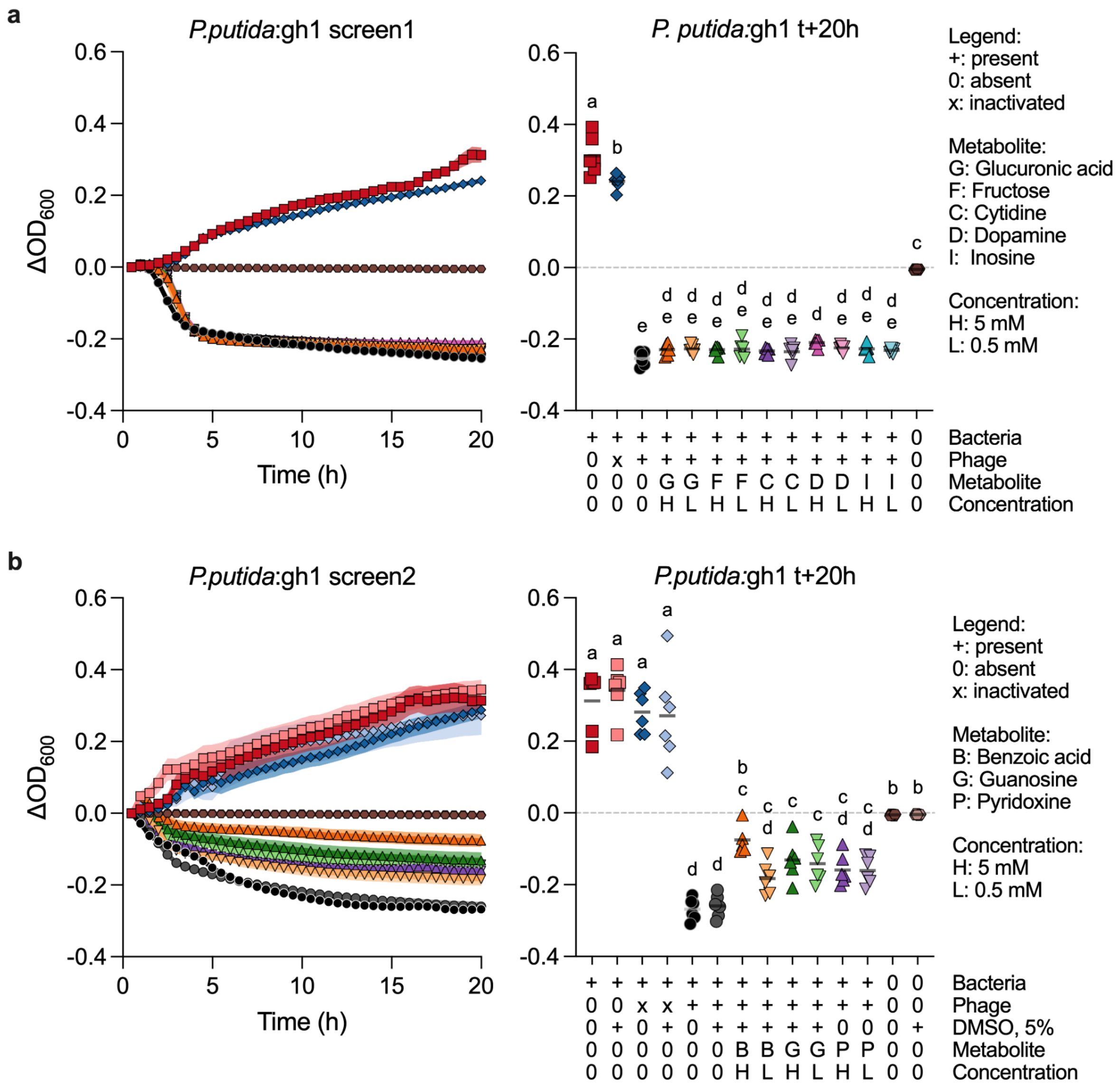


**Fig. S8: Initial bioassays testing exometabolites effects on phage infection *in vitro*.** *P. putida* A.3.12 cultures ($OD_{600}$ ~0.5 in LB medium) were supplemented with compounds (0.5 or 5 mM) and infected with phage gh-1 (MOI 0.01, $V$=200 µl). The $\Delta OD_{600}$ was measured for 20 h and compared to 0 mM metabolite controls: infected cells (black circles), uninfected cells (red squares), inactivated phage (autoclaving, blue diamonds), and uninoculated LB medium (brown hexagons). (**a**) Bioassay 1: Glucuronic acid, Fructose, Cytidine, Dopamine, and inosine. (**b**) Bioassay 2: Pyridoxine, Benzoic acid (+DMSO), Guanosine (+DMSO), and corresponding DMSO controls. Growth curves show an average $\Delta OD_{600}$ ± SE. Scatter plots show all values and mean (horizontal line) at t+20h. Different letters show statistical differences at $p<0.05$, One-way ANOVA with Tukey test, $n$=6 (separate mixtures/infections).

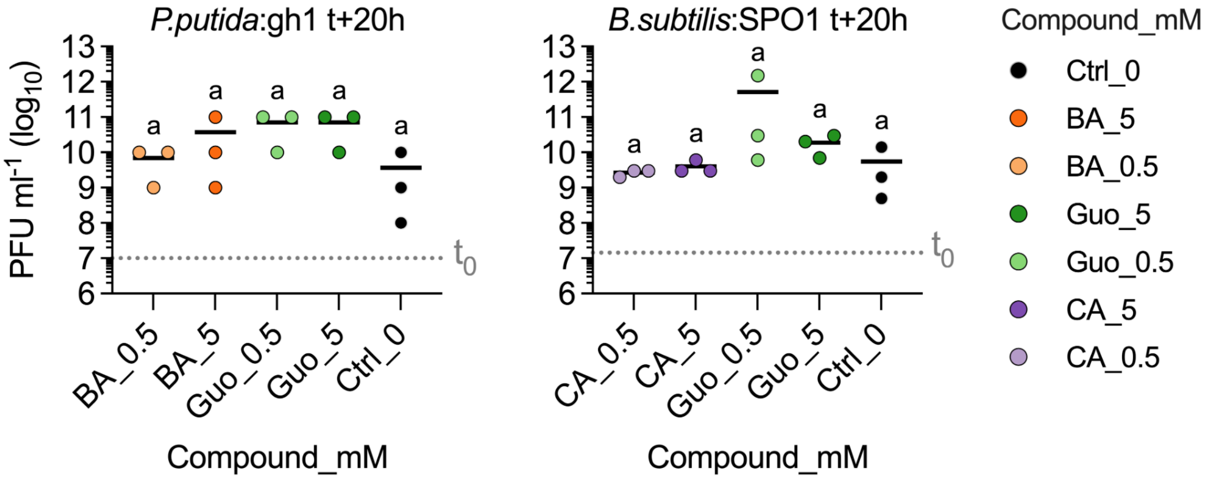


**Fig. S9: Effect of exometabolites on phage titer.** The bioassays determined phage concentrations by plaque assays in randomized 3 replicates of infected cultures in selected metabolite treatments after 20 h from phage addition. The horizontal black lines indicate the mean phage titer at the end of the experiment (t+20h). The host cells were infected at the MOI of 0.01. The horizontal dotted gray lines indicate the theoretical phage titer at the start of the experiment ($t_0$). Different letters indicate significant differences at $p$<0.05, One-way ANOVA with Tukey's test, $n$=3 (separate infections).
